# Hemagglutination by *B. fragilis* is mediated by capsular polysaccharides and is influenced by host ABO blood type

**DOI:** 10.1101/2020.08.19.258442

**Authors:** Kathleen L. Arnolds, Nancy Moreno-Huizar, Maggie A. Stanislawski, Brent Palmer, Catherine Lozupone

## Abstract

Bacterial hemagglutination of red blood cells (RBCs) is mediated by interactions between bacterial cell components and RBC envelope glycans that vary across individuals by ABO blood type. ABO glycans are also expressed on intestinal epithelial cells and in most individuals secreted into the intestinal mucosa, indicating that hemagglutination by bacteria may be informative about bacteria-host interactions in the intestine. *Bacteroides fragilis*, a prominent member of the human gut microbiota, can hemagglutinate RBCs by an unknown mechanism. Using a novel technology for quantifying bacterial hemagglutination, genetic knockout strains of *B. fragilis* and blocking antiserums, we demonstrate that the capsular polysaccharides of *B. fragilis*, polysaccharide B (PSB), and PSC are both strong hemagglutinins. Furthermore, the capacity of *B. fragilis* to hemagglutinate was much stronger in individuals with Type O blood compared to Types A and B, an adaptation that could impact the capacity of *B. fragilis* to colonize and thrive in the host.

**Importance Statement:** This study found that the human pathobiont, *B*. *fragilis*, hemagglutinates human red blood cells using specific capsular polysaccharides (PSB and PSC) which are known to be important for interacting with and influencing host immune responses. Because the factors found on red blood cells are also abundantly expressed on other tissues and in the mucous, the ability to hemagglutinate sheds light on interactions between bacteria and host throughout the body. Intriguingly, the strength of hemagglutination varied based on the ABO blood type of the host, a finding which could have implications for understanding if an individual’s blood type may influence interactions with *B*. *fragilis* and its potential as a pathogen versus a commensal.

## Introduction

The direct agglutination of red blood cells (RBC) (hemagglutination) is an important facet of bacterial physiology and has been shown to correlate with bacterial adherence and colonization of the host ^1^. While the antigens that determine blood type are canonically associated with RBCs, they are widely expressed by a range of other tissues, including intestinal epithelial cells and in approximately 80% of individuals – commonly referred to as “secretors”, blood glycans are secreted in high abundance at mucosal surfaces, including in the intestine ^2 3 4^. Therefore, hemagglutination can be indicative of a variety of aspects of bacteria-host interaction.

Numerous studies have linked hemagglutination with the pathogenesis and virulence of bacterial infections ^5 6^, as direct binding to host cells allows for the delivery of toxins, enzymes, and immune modulatory factors facilitating bacterial manipulation of host responses ^7 8 9^. For instance, Escherichia *coli* strains that are able to hemagglutinate are more virulent, in part due to their ability to adhere to host epithelial cells and initiate biofilm formation ^6^. Bacterial hemagglutination of RBCs is mediated by interactions between bacterial cell components and RBC envelope glycans that vary across individuals by ABO blood type and can be the basis of observed blood type specificity to infectious disease ^10 4^. For instance, sialic acid binding hemagglutinins produced by *Helicobacter pylori* play an important role in colonization of the intestinal mucosa and host blood type can influence the ability of *H. pylori* to target these receptors ^11 12^.

While the majority of studies on hemagglutination have focused on pathogens, the benefits conferred by this capability could extend to commensal bacteria, as the need to tightly adhere in the gastrointestinal tract could affect niche acquisition of gut microbiota in this densely populated environment ^13 14 15^. Some bacteria can also utilize host glycans as a food source, suggesting that hemagglutination may also represent a nutrient foraging strategy for bacteria, thus this phenotype may play a role in the establishment of spatial and metabolic niches^16^.

*Bacteroides fragilis* is a common component of the healthy human microbiome with capacity to protect the host from inflammatory diseases ^17^. However, *B*. *fragilis* also has pathogenic potential, as it can cause intraabdominal abscesses and enterotoxigenic strains can cause diarrhea^18 19^. Numerous reports have noted the ability of *B*. *fragilis* to hemagglutinate and strains of *B*. *fragilis* with higher hemagglutinating activity were found to be the most adhesive to human cells lines and were more frequently isolated from clinical specimens than from healthy fecal donors^27^. Experiments exploring the molecular factor(s) responsible for hemagglutination have indicated their presence on the bacterial capsule as well as in outer membrane vesicles (OMVs) but the specific factor driving this phenotype remains to be described^28^. However, efforts to characterize the class of molecule driving hemagglutination have suggested that this process is likely mediated by a polysaccharide as treatment with sodium periodate (which oxidizes polysaccharides) ablates hemagglutination, but treatment with proteinases or carbohydrates failed to inhibit hemagglutination^8^.

In total, *B*. *fragilis* is known to make eight capsular polysaccharides (CPSs) PSA-PSH. *B. fragilis’* interaction with the host immune response is driven in part by polysaccharide A (PSA), a zwitterionic polysaccharide (ZPS) characterized by repeating positive and negatively charged subunits^19^ that has broad effects including enhancing the ability of *B*. *fragilis* to colonize the host ^20 21^. PSB is the only other zwitterionic polysaccharide and shares at least some of PSA’s capacity for immune-modulation and impacts on the host^34^. Of the eight CPSs, the expression of all but PSC are tightly controlled by a site-specific recombinase, Multiple promoter invertase (Mpi) which mediates the inversion of the promoter regions ^22 23^. The ability, not only to express diverse surface structures but to orchestrate their expression may allow commensal bacteria to quickly adapt to different environments they encounter in the host, evade immune responses, and utilize varied nutrient resources^24^. Among the 8 CPS, PSC is the only CPS whose expression is controlled independently^22^. These findings prompted us to hypothesize that the CPSs produced by *B*. *fragilis*, that have been shown to be important for adherence, colonization, and immune modification, may be the factors mediating hemagglutination by *B*. *fragilis*. Utilizing a novel technology to quantify bacterial hemagglutination by *B*. *fragilis*, genetic knockout strains of *B*. *fragilis* and blocking antiserums, we show that PSB and PSC are strong hemagglutinins. Furthermore, the capacity of *B*. *fragilis* to hemagglutinate was much stronger in individuals with Type O blood compared to Types A and B, an adaptation that could impact *B*. *fragilis* colonization of humans based on their blood type.

## Results

### Capsular polysaccharides PSB and PSC contribute to hemagglutination by B. fragilis

To explore hemagglutination in *B*. *fragilis*, we performed assays in which suspensions of RBCs from healthy adults were mixed with serial dilutions of bacteria and hemagglutination was imaged and quantified using a CypherOne plate reader (InDevR, Boulder Colorado). The CypherOne plate reader enables robust quantification of the strength of hemagglutination utilizing automated imaging and standardized measurements of hemagglutination. The plate reader assigns a non-agglutination score, that is inversely proportional to hemagglutination, to each well in 96 well plates containing RBCs and a dilution series of an antigen being tested (Fig. 1). This plate reader also makes a titer call which is the dilution at which the hemagglutination phenotype is lost (as indicated by the highlighted well) (Fig. 1B). The CyperOne plate reader was originally developed for influenza diagnostics and the evaluation of vaccine candidates; prior to this study it had not been used with bacteria.

**Figure 1.**
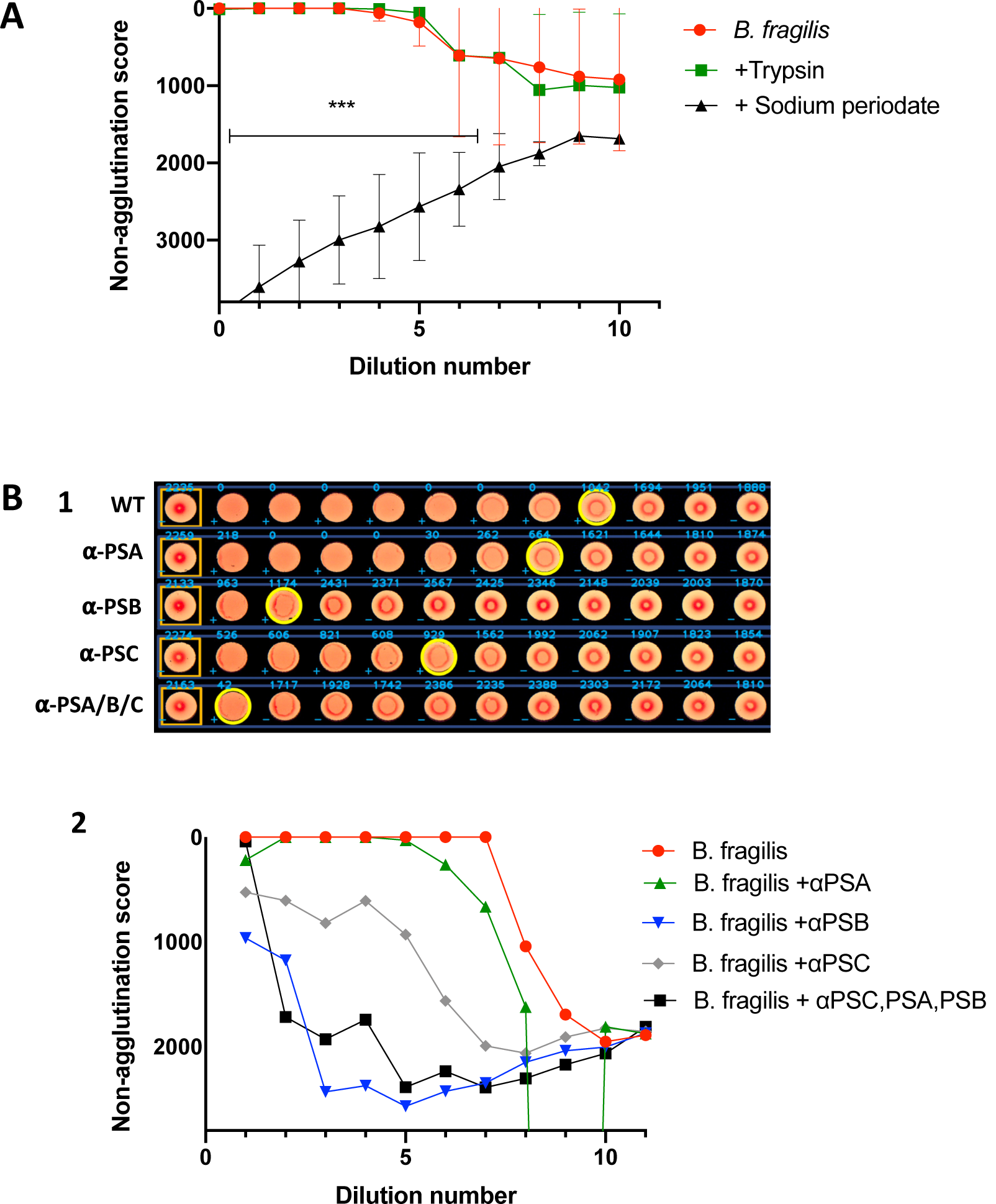
Disruption of all capsular polysaccharides or targeted blocking of PSB and PSC inhibits hemagglutination by *B*. *fragilis*. **A)** Hemagglutination patterns of untreated *B. fragilis, B. fragilis* that had been pre-incubated with trypsin, or with sodium periodate were compared using RBC from 3 individuals. *** indicates a *p* = < 0.0001 as determined by Welch’s T-test. **B). 1**. Plate analysis of *B*. *fragilis* WT versus *B*. *fragilis* WT that had been incubated with antiserum to capsular polysaccharides. RBCs from healthy human donors were mixed with an equal volume of either PBS (first column) or 1.0 × 10^8^ cfu/ml of washed *B*. *fragilis* WT or *B*. *fragilis* + antiserums. Material in each well was than serially diluted 3:2 down the row. The plate was imaged and analyzed with a Cypher One™ Instrument, 4.0 Analysis Software. Non-agglutination results in a button of RBC at the bottom of the well as seen in column 1 with the PBS negative control. Complete agglutination results in a consistent suspension of RBC as seen in column 2 with untreated WT *B*. *fragilis*. Non-agglutination scores are assigned to each well by the software (blue text) and are inversely proportional to hemagglutination, with complete agglutination having a score of 0. The wells highlighted in yellow indicate the dilution at which hemagglutination is lost as determined by the software. **2**. Plot of non-agglutination scores for the imaged plate. Welch’s t-test was used to assess significance between *B*. *fragilis* WT and antibody treated strains (WT vs *α*PSA *p* = N.S., WT vs *α*PSB *p* = 0.0002, WT vs *α*PSC *p* = 0.0339, WT vs *αPSA/B/C*= 0.0009).

We first performed this hemagglutination assay to compare wild-type (WT) *B*. *fragilis* NCTC 9343 to bacterial suspensions that had been treated with either sodium periodate to destroy polysaccharides or trypsin to destroy proteins for 60 minutes (Fig. 1A). *B*. *fragilis* WT treated with sodium periodate ablated the hemagglutination phenotype while treatment with trypsin had no effect, supporting previous reports that hemagglutination by *B*. *fragilis* was mediated by polysaccharide (Fig. 1A)^8^. We next evaluated hemagglutination in WT *B*. *fragilis* that had been incubated with antiserums to PSA, PSB or PSC or with mixture of all three (Fig. 1B). While blocking with each individual antiserum resulted in a partial loss of phenotype, blocking of PSB or PSC having a more profound impact than blocking of PSA, simultaneous blocking of all three CPS resulted in a full loss of the hemagglutination phenotype (Fig. 1B). The impact of each CPS of interest was further assessed by comparing hemagglutination by WT *B*. *fragilis* NCTC 9343 to isogenic knockouts of PSA, PSB, or PSC operons (*B*. *fragilis* ΔPSA, ΔPSB, or ΔPSC) ^26^ (Fig. 2A; Supplemental Fig. 1). We also qualitatively observed hemagglutination by light microscopy (Fig. 2B). We used the titer call of the loss of hemagglutination phenotype to plot survival curves (Fig. 2C) and to estimate hazard ratios for each strain (Fig. 2D). While all three mutants showed a similar pattern of increased risk of phenotype loss, the effect was only statistically significant for ΔPSB and ΔPSC, with the latter showing the strongest effect.

**Figure 2.**
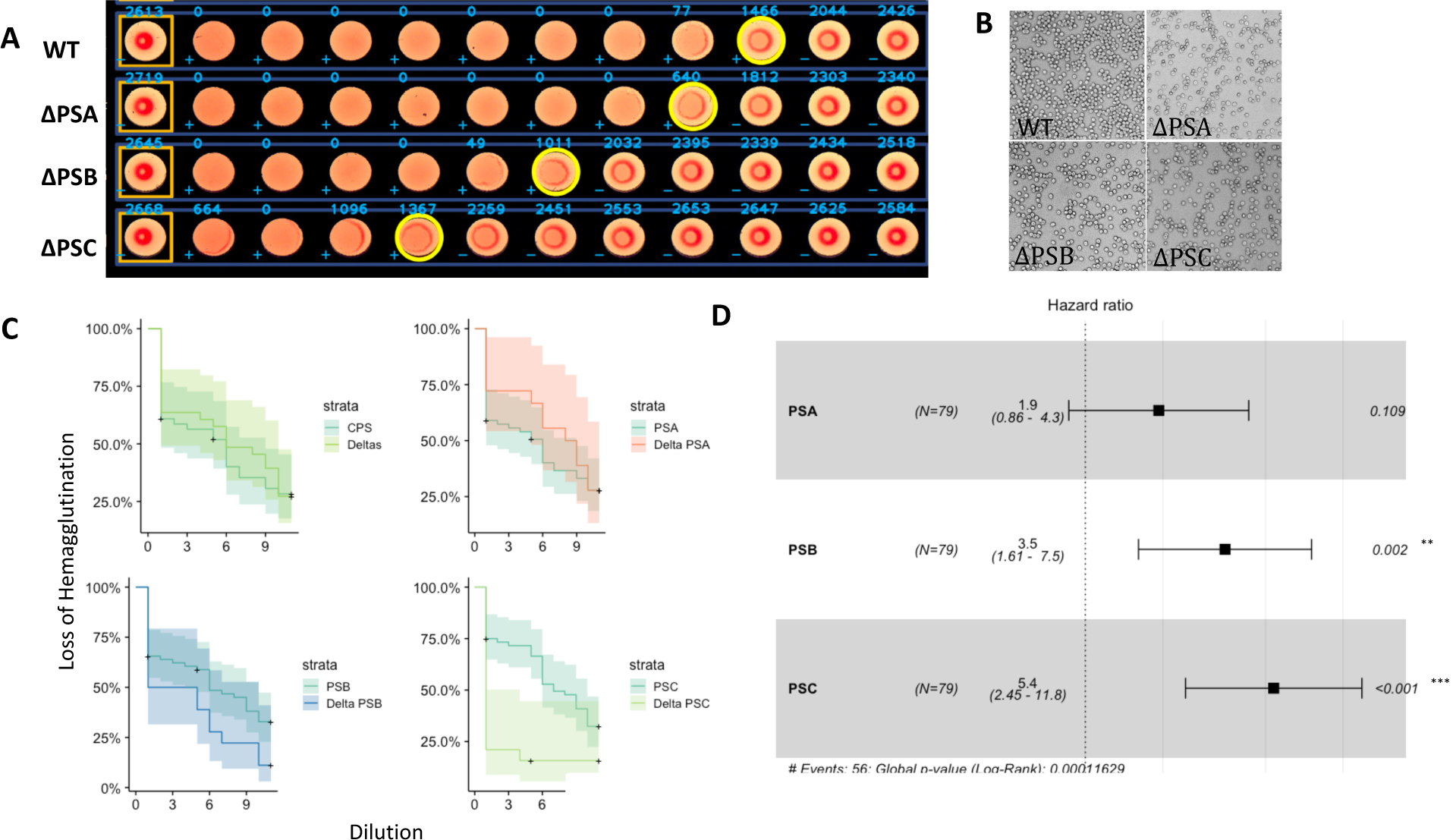
Capsular polysaccharides PSB and PSC contribute to hemagglutination by *B*. *fragilis*. **A**. RBCs from healthy human donors (Type O) were mixed with either PBS (first column) or 1.0 × 10^8^ cfu/ml of *B*. *fragilis* WT or delta strains (second column). Material in each cell was than serially diluted 3:2 down the row. The plate was imaged and analyzed with a Cypher One™ Instrument, 4.0 Analysis Software. *B*. *fragilis* WT versus isogenic knock-out strains. **B**. Light microscopy of hemagglutination patterns of human RBC at 20x magnification mixed with 1.0 × 10^8^ cfu/ml *B*. *fragilis* WT, *B*. *fragilis* ΔPSA, B. *fragilis* ΔPSB, or *B*. *fragilis* ΔPSC. **C**. Plots of dilution to loss of phenotype for *B*. *fragilis* (clockwise) WT versus any delta, any strain with PSA vs those without, strains with PSB vs those without, strains with PSC vs those without analyzed with the Cox proportional-hazards model (n=26). **D**. Hazard ratios of the risk of loss of phenotype for each mutant strain compared to strains expressing that CPS.

### Expression of the PSB and PSC can compensate for the deletion of PSA

Although all knock-out strains showed an overall reduction in hemagglutination, *B*. *fragilis* ΔPSA had a smaller reduction in hemagglutination compared to ΔPSB and ΔPSC that did not significantly differ from *B*. *fragilis* WT. It has been previously reported that the deletion of one CPS operon can result in increased expression of another^22^. To address whether PSA is just the weakest hemagglutinin or if smaller reduction in hemagglutination in ΔPSA was driven by compensatory expression of PSB or PSC, we used reverse transcriptase polymerase chain reactions (RT-PCR) to compare expression levels of the three CPS between WT and mutant strains. *B*. *fragilis* ΔPSA had increased expression of PSB and PSC compared to *B*. *fragilis* WT (Supplemental Fig. 2, however the extent to which expression was increased ranged widely between biological replicates indicating stochasticity (F-test of equality of variances *p= 0.0044* PSB, *0.00054* PSC). This indicates that the lack of a strong loss of the hemagglutination phenotype for ΔPSA could be influenced by compensation by PSB and PSC. However, we note that the blocking antiserum to PSA also had a weak phenotype (Fig. 1B), suggesting again that PSB and PSC are more important hemagglutinins.

### The extent of the hemagglutination phenotype varies based on ABO blood type

Since bacterial hemagglutination of RBCs can be mediated by interactions between bacterial cell components and RBC envelope glycans that vary across individuals by ABO blood type, we explored the relationship between ABO blood type and strength of hemagglutination in our assays, which were performed on RBCs from 26 healthy adults; 14 individuals with type O blood, 7 with type A, and 5 with type B (Fig. 3, Supplemental Fig.3). The hemagglutination phenotype was the strongest with type O blood and was profoundly reduced with type A or B blood (Fig. 3A). We modelled the relationship between serial dilution and the loss of hemagglutination phenotype using survival models by blood type (Fig 3B) and found elevated risk of phenotype loss among “non-O” blood type (type A or B) relative to O (HR=9.4; CI:2.5-36), indicating a marked affinity for type O blood over types A or B (Fig. 3B). Analysis by 2-way ANOVA indicated that there was not a significant interaction between blood type and bacterial strain. The influence of bacterial strain accounted for 20.57% (P-value < 0.0001) of the observed variation, host blood type for 15.53% of the variation (P-value < 0.0431), and intrasubject variation accounted for a further 34.70% (P-value < 0.0001).

**Figure 3.**
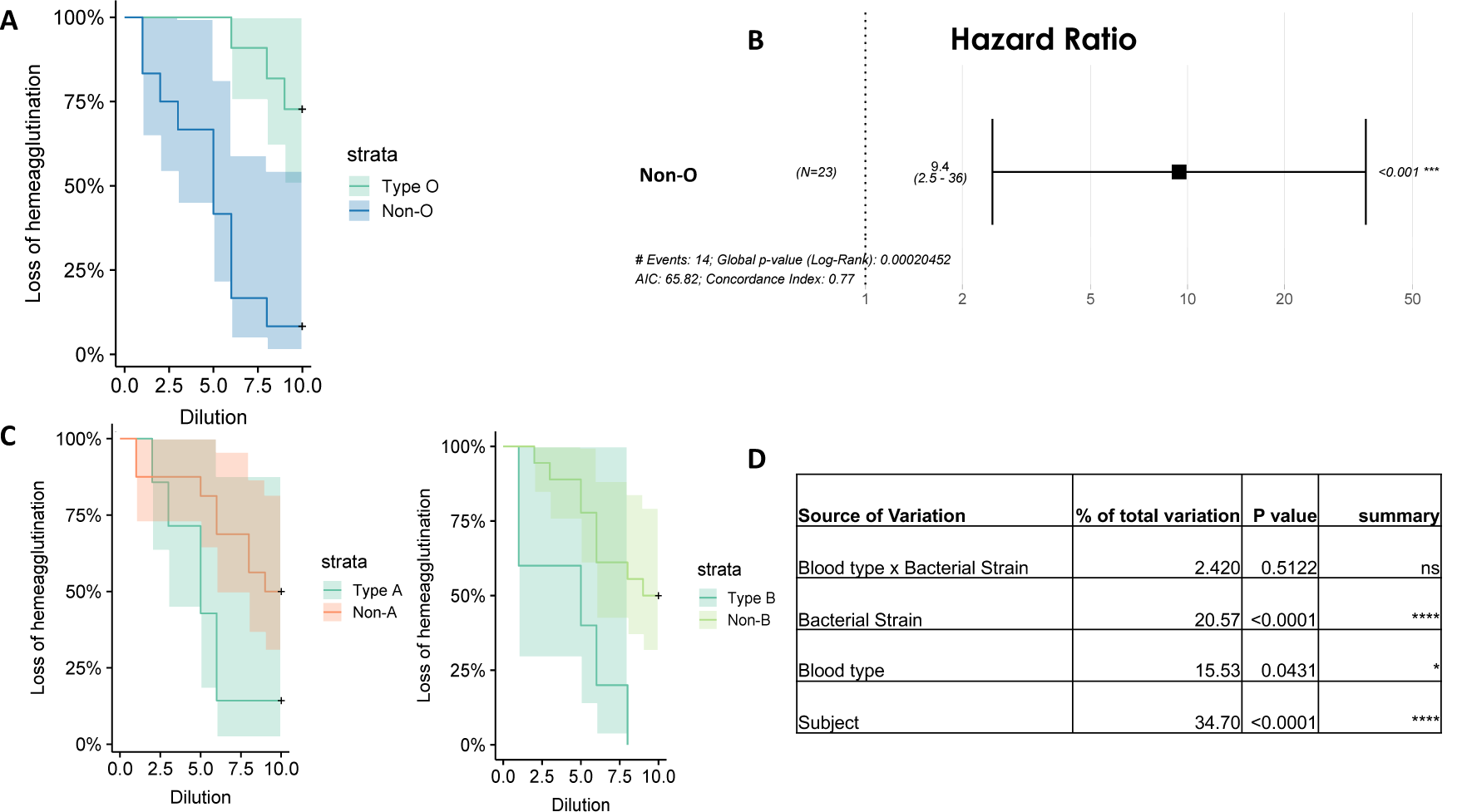
The degree of the hemagglutination phenotype varies based on ABO blood type. **A**. Plots of dilution to loss of phenotype for *B*. *fragilis* WT with Type O blood versus non-O blood (Type A or B) analyzed with the Cox proportional-hazards model. **B**. Hazard ratios (HR 9.4; CI 2.5-36) for non-O blood (type A or AB) as compared to Type O. **C**. Hazard ratios for being Type A, or Type B. **D**. 2-way ANOVA analysis of influence and interaction of blood type and bacterial strain and variance by subject.

## Discussion

It has been over thirty years since it was first reported that *B*. *fragilis* is able to hemagglutinate RBCs ^29^ and nearly twenty since it was determined that this capacity is driven by a component of the bacterial capsule ^8^. Here we show that interfering with either PSB or PSC, either by blockade of the CPS with specific antiserum or with genetic knock-out strains, reduces the overall capacity of *B*. *fragilis* to hemagglutinate. Although, inhibition of PSA showed some reduction in the hemagglutination phenotype, the effect was highly variable and did not achieve statistical significance. Blocking PSA, PSB, and PSC simultaneously, completely ablates hemagglutination.

While there is much to learn about the specific roles of the different CPS of *B*. *fragilis*, it is clear that they are important to the bacteria as efforts to maintain a strain that completely lacks the CPS repertoire have failed and strains expressing only a single CPS were unable to effectively colonize the host^22 25^. The functional activities of PSA, a CPS that unlike most has a zwitterionic structure, have been extensively studied and include the ability to induce IL-10 expression by regulatory T-cells^30^, to attenuate inflammation in models of Inflammatory Bowel Disease (IBD)^31^, and to cause intra-abdominal abscess^32^. PSA also influences the ability of *B*. *fragilis* to colonize the intestinal mucosa, as *B*. *fragilis* ΔPSA showed a mucus-specific colonization defect in mice ^33^. PSB and PSC are less studied, but there is evidence that they similarly play important roles in colonization by *B*. *fragilis*^22^. PSB is also zwitterionic and shares at least some of PSA’s capacity for immune-modulation impacts on the host^34^.

Out of the eight CPSs (PSA-PSH) produced by *B*. *fragilis*, seven are phase variable and their expression can be controlled by invertible promoters that are regulated by the recombinase, multiple promoter invertase (mpi)^23^. PSC alone is regulated via a non-invertible promoter, which has been purposed to serve as a safety net, ensuring that *B*. *fragilis* always has at least one CPS expressed ^25^. The CPS repertoire of *B*. *fragilis* is highly flexible and redundant and disruption of one capsular polysaccharide can be compensated for by the induction of expression of another of the eight distinct CPS components^26^. Our results further support this finding, as gene transcript analysis by RT-PCR indicated that the loss of PSA can result in an increase in the expression of the PSB and PSC.

The interplay between PSB and PSC, as well as their importance to *B. fragilis’* ability to colonize the mammalian host, is highlighted by an elegant series of experiments by Liu et. al ^22^. Using knockout strains of the 8 different CPS, they demonstrated a marked increase in PSC expression when PSB alone was knocked out. If all of the other 7 CPS were disrupted, PSC, which is only expressed at low levels in WT *B*. *fragilis*, was expressed by 100% of cells. These findings support the hypothesis that *B*. *fragilis* is reliant on the expression of at least one CPS with PSC acting as the backup. Interestingly, when they also shut down PSC in the strain lacking the other 7 CPS, an alternate recombinase was able to rescue expression of PSB^22^. Strains of *B*. *fragilis* lacking any CPS are unable to colonize in a mouse model, and while a single CPS can facilitate colonization, these strains cannot outcompete wild-type *B*. *fragilis* with a complete CPS repertoire ^22 25.^ This previous work supports the hypothesis that PSB and PSC represent a tightly coordinated and redundant system that would ensure maintenance of the hemagglutination phenotype. Indeed, the ability to hemagglutinate may serve as a mechanism for bacterial colonization of the mucosa or facilitate vesical delivery allowing these molecules to gain the proximity necessary to exert their influence on host cells. It is attractive to speculate that these factors may work in concert-with hemagglutination by PSB and PSC facilitating proximity to host where PSA can exert its influence on host responses.

Hemagglutination can benefit bacteria in multiple ways, including facilitating pathogenicity, adhering to host cells, and utilizing host glycans as a food source ^6 35^. Targeting blood antigens both as a means to adhere to the host and as a nutrient resource is a well-known strategy of many pathogenic bacteria^10^. The blood glycans to which bacteria typically bind during hemagglutination are present on many cells throughout the body, including on epithelial cells in the intestinal lining, suggesting that hemagglutination has the potential to facilitate *B*. *fragilis* colonization. However, about 80% percent of individuals are also characterized as “secretors,” meaning that blood glycans are secreted at large amounts in bodily fluids, including into the intestine^36 10^. It has been proposed that this phenotype allows individuals to clear potential pathogens that hemagglutinate from the gut and other body sites^37^. Interestingly, host blood type can influence susceptibility to a number of pathogens indicating that some bacteria may target specific blood antigens^12 4 38^. Intriguingly, our results suggest that the hemagglutination capacity of *B*. *fragilis* varies based on the host’s ABO blood type, with agglutination of type A and B RBC by *B*. *fragilis* being significantly reduced compared to hemagglutination of type O blood. While type O blood consistently hemagglutinated to a greater extent than non-O, we observed significant inter-individual variation in hemagglutination capacity within blood types which may reflect the variability of ABO expression levels on RBC^39 40^. We didn’t observe any difference based on Rh status, but that could be the result of only using a small number (n=5, 4 O-, 1 A-) of Rh-negative blood samples in our assays.

While there are some compelling studies suggesting that host ABO blood type might influence the composition of the microbiome^41 42 43^, others have not found associations between the microbiome and host blood type ^44^. Our finding that a common and influential pathobiont, *B*. *fragilis*, has an increased hemagglutination capacity for host blood type O versus non-O *in-vitro* supports that some commensal bacteria have blood type specificity as has been described for pathogens. It would be valuable to assess the ability of *B*. *fragilis* to adhere to epithelial cells expressing different blood glycans. These findings warrant further investigation to determine if host blood type influences the presence or abundance of *B*. *fragilis* as a resident in the microbiota, or if blood type is a risk factor for abscess caused by *B*. *fragilis*. These findings justify including blood type in microbiome analyses, which could have important implications for analyzing the composition of individual microbiomes, and for informing microbial prevention or treatment strategies.

## Methods

### Preparation of RBC

Human red blood cells were harvested from healthy donors by a blood draw (COMIRB No: 17-0348, 19-2556) followed by Ficoll gradient isolation. Cells were then pelleted at 1500 rpm for 10 minutes, washed three times with 1X PBS, and resuspended at a concentration of 1% in PBS. RBC were added to the bacterial suspensions and the plates were covered and incubated for 60 minutes at 4°C. Plates were then imaged and hemagglutination was quantified using the Cypher One™ Instrument and 4.0 Analysis Software (InDevR, Boulder, CO).

### Bacterial strains and associated reagents

WT *B*. *fragilis* (NCTC 9343) was purchased from ATCC. *B*. *fragilis* Δ PSA, *B*. *fragilis* Δ PSB, and *B*. *fragilis* Δ PSC, as well as the polyclonal rabbit antiserums, were generously shared with our lab by Dr. Sarkis Mazmanian (Caltech) and Dr. Gregory Donaldson (Rockefeller University). The methods used to produce and validate these knock-out strains have been previously described ^26 45^. All bacterial isolates were grown in rich media (Mega Media) prepared in our lab ^46^.

### Hemagglutination Assays

Bacterial cultures were started out of glycerol and were grown overnight in liquid Mega Media to early stationary phase. Because *B*. *fragilis* cultures can be heterogenous in capsule phenotype, we normalized cultures using Percoll density centrifugation as previously described ^47^. The fraction found to hemagglutinate most effectively for all strains was at the 20-40% interface, this fraction was passed once to remove Percoll and grown overnight in liquid Mega Media to early stationary phase. CFU/mL were determined by staining aliquots of 18-hour cultures with the BD Biosciences Bacterial Detection Live/Dead kit and enumerating live bacteria by flow cytometry. Bacterial cultures were then normalized to 1.0 × 10^8^ cfu/mL, pelleted at 2500 rpm for 10 minutes at room temperature, and washed twice with 1X PBS. The washed bacteria were then added to the second column of each row of a 96 well plate in equal volume to the human RBCs. We then serially diluted (1.5 dilution factor) with 1X PBS down the row, the first well of each row only got 1X PBS and served as a negative control.

### Statistical Analysis

We fit Cox proportional hazard models to estimate the relationship between serial dilution and loss of hemagglutination phenotype overall and stratified by A/B versus O blood type using R version 3.4.0^48^. Prism 8.3.1 was used to compare pellet density scores and to perform 2-way ANOVA to assess contributions to variation.

### Determining ABO blood type

ABO blood type was determined using Anti-A, Anti-B, and Anti-Rh(D) monoclonal murine antibodies (ThermoFisher Science). Agglutination indicated presence of blood antigens and was read by eye to determine donor blood type. The accuracy of this method was verified on a subset of samples that were also back typed using serum. Typing was done with the inclusion of positive controls and were read independently by two people.

### Blocking of PSA, PSB, and PSC with polyclonal antiserum

*B. fragilis WT* was pre-incubated with 1.5ng/ml in the first dilution for 30 minutes at 37°C. Bacteria that had been pre-incubated with the antiserum were then serially diluted and mixed with RBC, hemagglutination was imaged and quantified. To block PSA, PSB, and PSC simultaneously, *B*. *fragilis* WT was pre-incubated with 0.5ng/ml of each antiserum and analyzed as above.

### Microscopy

Bacteria/RBC suspensions (equal parts 1.0 × 10^8^ cfu/mL washed bacteria to 1% RBC) were spotted onto glass slides mounted with a coverslip and immediately imaged on a Zeiss Axioplan 2, 3i camera, and Slidebook6 software at a magnification of 20x.

### RT-PCR of CPS expression

RT-PCR was performed on extracted RNA from bacterial cultures used for the hemagglutination assays. RNA was extracted using the PureLink™ RNA Mini Kit (ThermoFisher), cDNA was synthesized using the RT^2^ First Strand Kit (Qiagen), the RTPCR was prepared using Kapa SYBR Fast qPCR master mix (Roche Wilmington, MA) and completed on the CFX96 platform (BioRad Hercules, CA). Relative expression levels of the CPS were compared to 16S rRNA and analyzed using the ΔΔCq method. Results are from three biological replicates and all samples were run with technical triplicates. Sequences of the primers that we used are shown in Supplemental Table 1.

## Acknowledgements

We would like to thank Dr. Sarkis Mazmanian and Dr. Gregory Donaldson for sharing the *B*. *fragilis* knock-out strains, PSA, PSB, and PSC anti-serums. We would also like to thank Dr. Gregory Donaldson for thoughtful input on this project. We want to express our gratitude to Garret Wilson at InDevr for his guidance in designing the hemagglutination assays and analyzing the data. We are grateful for input on data visualization from Janet Siebert and Casey Martin. Finally, we would like to thank Dr. Linda van Dyk for her support and insight on this project.

## Supplementary Materials

**Supplemental Figure 1.**
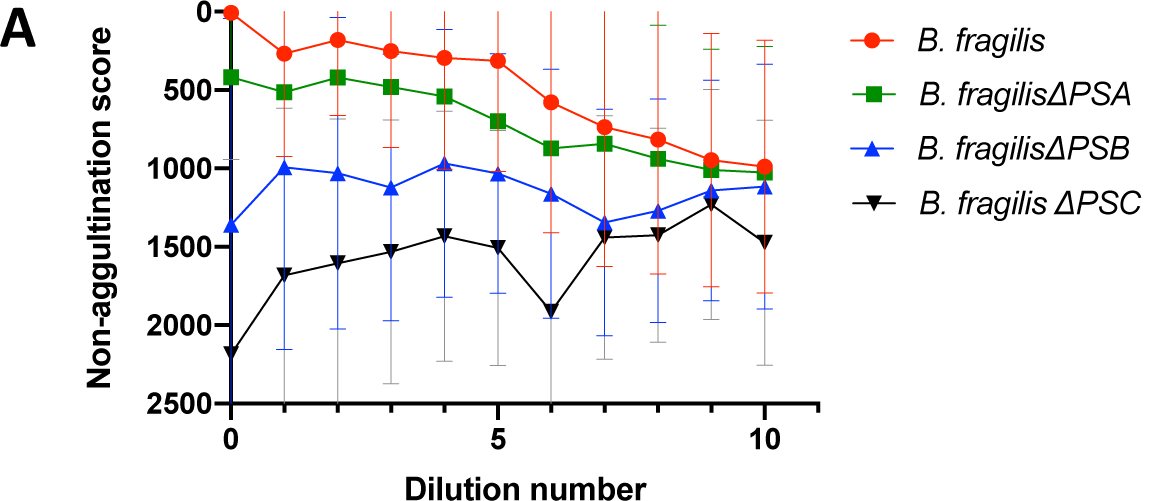
Capsular polysaccharides PSA, PSB, and PSC all contribute to hemagglutination by *B*. *fragilis*. **A**. Hemagglutination by *B*. *fragilis* WT was compared to the knockout strains, the plate was imaged and analyzed with a Cypher One™ Instrument, 4.0 Analysis Software. A non-agglutination score (which is inversely proportional to hemagglutination) is assigned to each well. Results are representative of 21 experiments on unique RBC (O=9, A=7, B=5). Welch’s t-test was used to asses significance between *B*. *fragilis* WT and KO strains (WT vs ΔPSA *p* = N.S., WT vs ΔPSB *p* = 0.0001, WT vs ΔPSC *p* = 0.0001

**Supplemental Figure 2.**
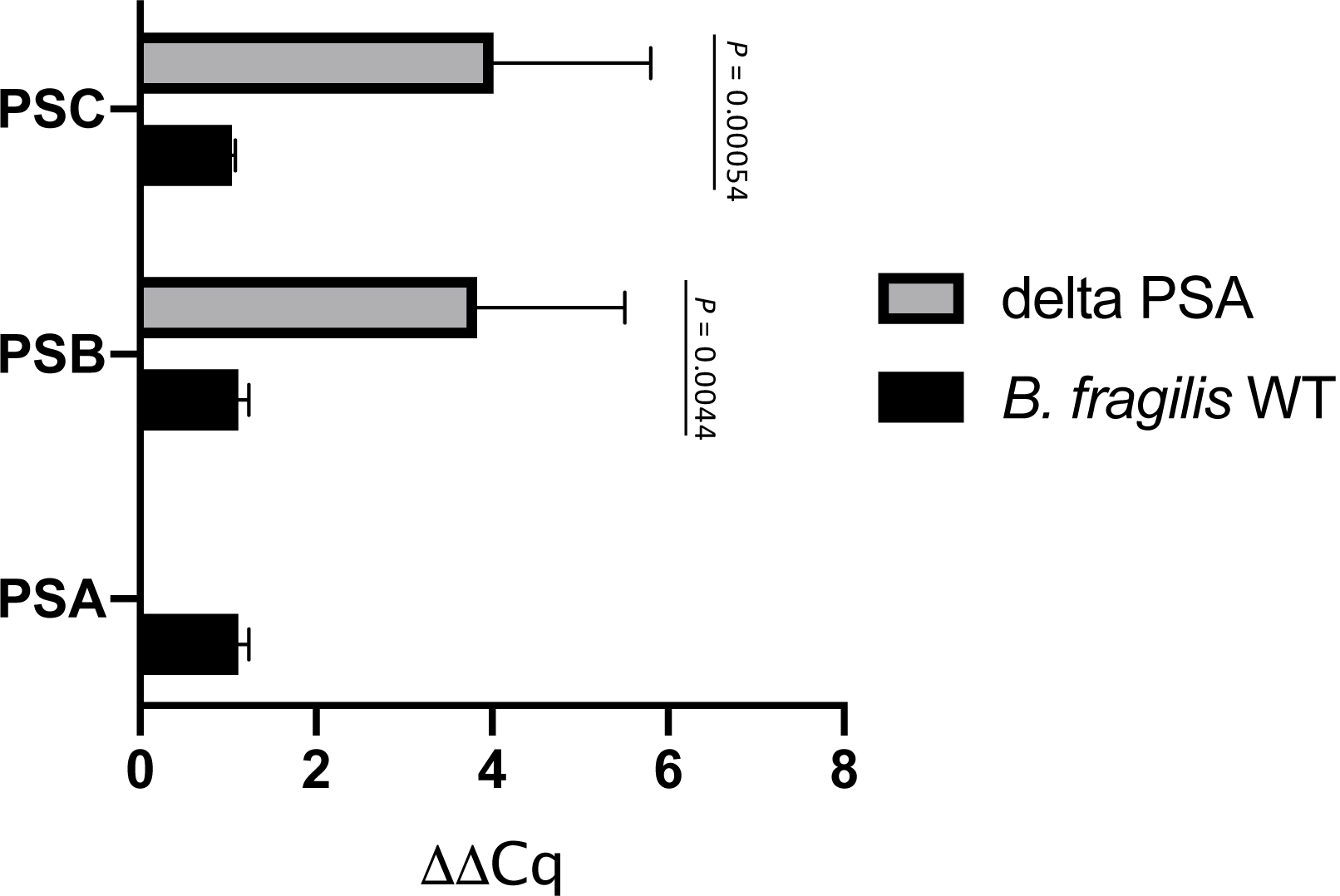
Expression of PSB and PSC are increased in ΔPSA. The relative normalized expression of PSA, PSB, and PSC were examined using RT-PCR. Expression of PSB and PSC is greater and highly variable in *B*. *fragilis* ΔPSA as analyzed by F-tests for equality of variance of relative PSB expression (*p= 0.0044)* and PSC expression (*p=* .*000054)*.

**Supplemental Figure 3.**
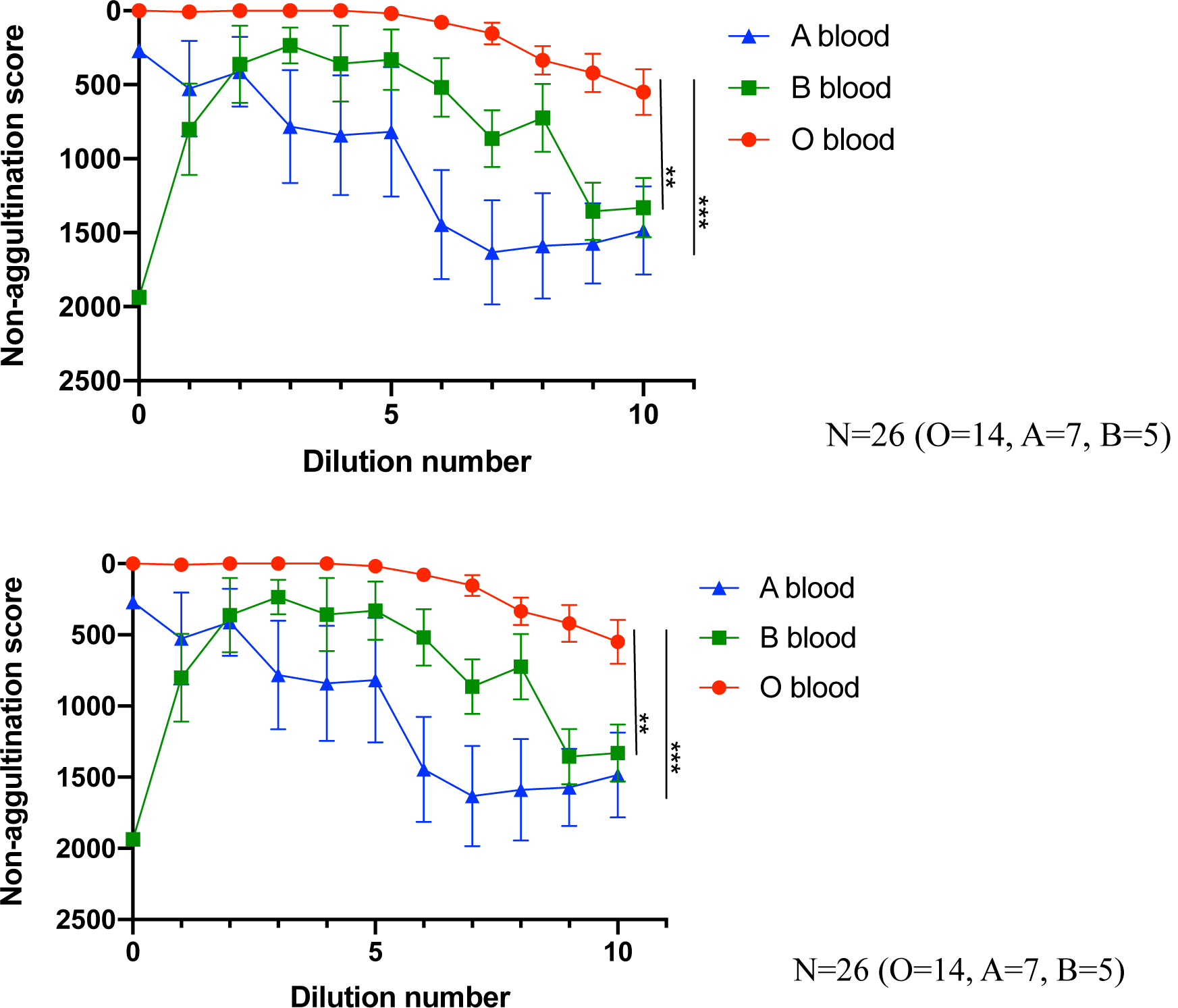
Hemagglutination by *B*. *fragilis* varies based on ABO blood type. Hemagglutination by WT *B*. *fragilis* on RBC from 26 individuals were compared based on ABO blood type (O=14, A =7, B=5). Significance of blood type as determined by Welch’s T-test is indicated. *** *p* = 0.0001 O vs. A, ** *p*= .0023 O vs. B. Non-agglutination scores (which are inversely proportional to hemagglutination) were assigned by analysis with a Cypher One™ Instrument, 4.0 Analysis Software.

**Supplemental Table 1.**
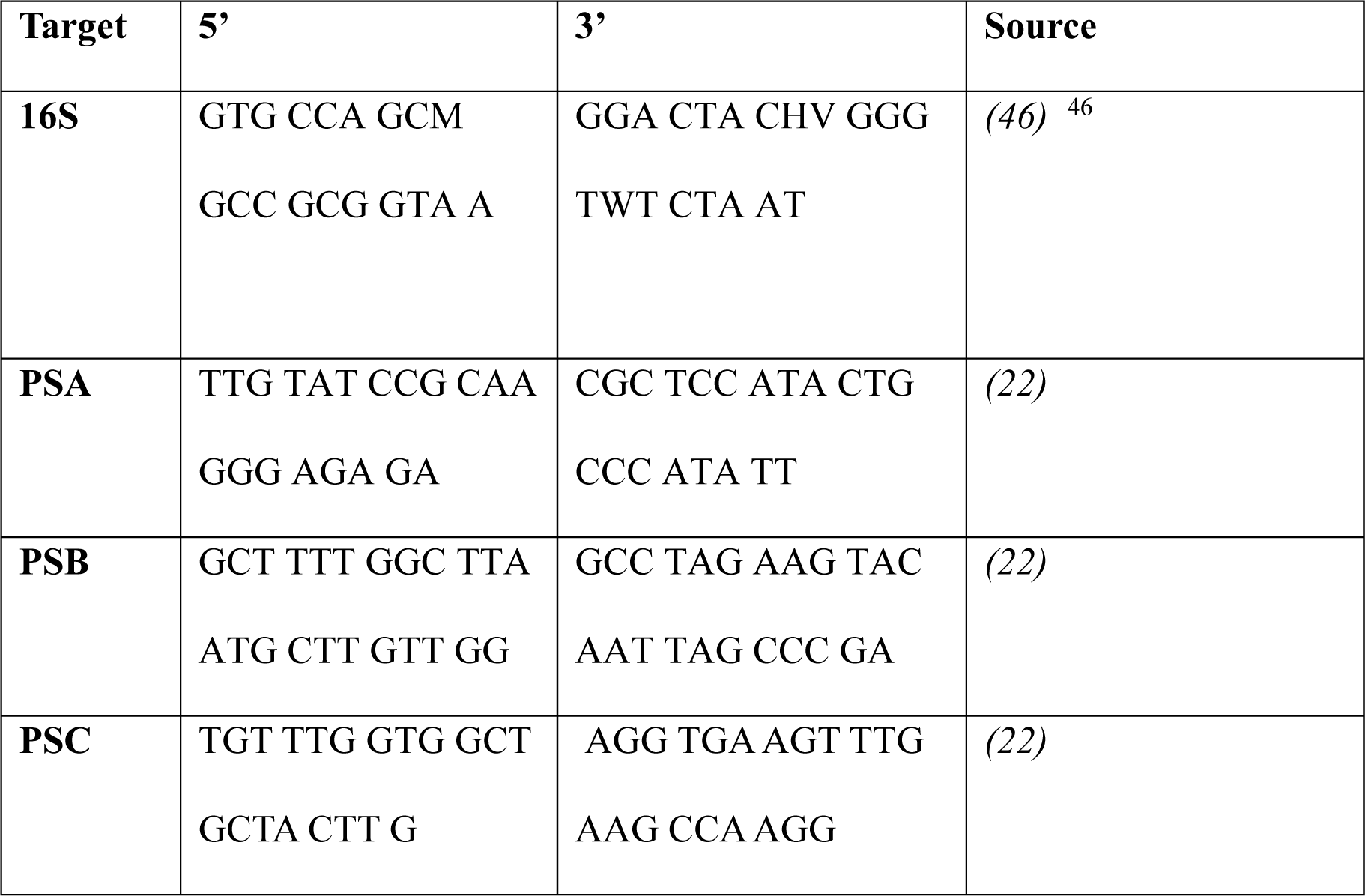
Primers used in this study.

